# Cancer mutational processes vary in their association with replication timing and chromatin accessibility

**DOI:** 10.1101/2021.05.05.442736

**Authors:** Adar Yaacov, Oriya Vardi, Britny Blumenfeld, Avraham Greenberg, Dashiell J. Massey, Amnon Koren, Sheera Adar, Itamar Simon, Shai Rosenberg

**Author notes:** **To whom correspondence should be addressed:** Rosenberg Shai, MD, PhD, Hadassah – Hebrew University Medical Center, Kiryat Hadassah, Jerusalem 91120, Israel, Itamar Simon, PhD, Department of Microbiology and Molecular Genetics, Institute of Medical Research Israel-Canada, Hebrew University Medical School, Hadassah Ein Kerem, Jerusalem 9112102, Israel. These authors contributed equally to this work (Adar Yaacov, Oriya Vardi). These authors co-supervised the research and manuscript (Itamar Simon, Shai Rosenberg).

## Abstract

**Background:** Cancer somatic mutations are the product of multiple mutational and repair processes, which are tightly associated with DNA replication. Distinctive patterns of somatic mutations accumulation in tumors, termed mutational signatures, are indicative of processes the tumors underwent. While tumor mutational load is correlated with late replicating regions and spatial genome organization, much is unknown about the association of many different mutational processes and replication timing, and the interplay with chromatin structure remains an open question.

**Methods:** We systematically analyzed the mutational landscape of 2,787 WGS tumors from 32 different tumor types separately for early and late replicating regions. We used sequence context normalization and chromatin data to account for sequence and chromatin accessibility differences between early and late replicating regions. Moreover, we expanded the signature analyses to doublet base substitutions and small insertions and deletions by developing an artificial genomes-based approach to account for sequence differences between various genomic regions.

**Results:** We revealed the replication timing (RT) association of single base, doublet base and small insertions and deletions mutational signatures. The association is signature specific: some are associated with early or late replication (such as UV-exposure signatures SBS7b and SBS7a, respectively) and others have no association. Most associations exist even after normalizing for genome accessibility. We further developed a focused mutational signature identification approach, which uses RT information to improve signature identification, and found that SBS16, which is biased towards early replication, is strongly associated with better survival rates in liver cancer.

**Conclusions:** Our comprehensive analyses enabled a more robust classification of RT association of single base, doublet base and indels signatures. By doing so, we demonstrated a variation in the association with RT, as many mutational processes biased towards either early or late replication timing, and others have an equal RT distribution. These associations were independent from chromatin accessibility in most cases. This work highlights that restricting signatures analyses to concise genomic regions improves identification of signatures, such as SBS16, and demonstrates its clinically relevance as a predictor of improved survival of liver cancer patients.

## Introduction

Somatic mutations in cancer genomes are accumulated along all stages of the cell lineage and are the summation of multiple mutational processes^1^. Different mutational processes generate unique combinations of mutation types, termed “Mutational Signatures”. Systematic analysis of the frequency of somatic mutations in its immediate genomic context is indicative of the mutational processes that were active in the tumor cells^2,3^. Such analysis of mutations frequency revealed many mutational signatures that are indicative of various mutagenesis processes^4^. Some of the signatures (such as single base substitution (SBS) signature 1) are found in all tumor types, indicating that they stem from a very general mutagenesis process, whereas others (such as SBS7a-d) are characteristic of a single type of cancer due to a tissue-specific mutagenesis process (in this case UV damage in skin cancer). In recent years, analyses of mutational signatures were expanded from single base substitutions (SBS) to include also doublet base substitutions (DBS) and small insertions and deletions (indels)^5^.

The DNA replication procedure plays an important role in mutagenesis^6^, as mismatches can be introduced and DNA damage may be fixed into mutations. Indeed, several replication features (such as fork rate and direction) are known to be associated with certain types of mutation loads^7,8^. Moreover, replication timing (herein called RT), the relative time in S phase that each genomic region is replicated^9^, is found to be associated with mutation load. The RT of a region reflects a higher order of genomic organization as it correlates with basic chromosomal features such as the regional GC content, Giemsa banding and gene density. In addition, early replicating regions are packed in more accessible chromatin and are more involved in transcription^10^. RT is strongly associated with mutation rates of both germ line and somatic mutations, which are much higher in genomic regions that replicate later in S phase (reviewed in^11^), suggesting that either mutagenesis or repair occurs in different intensities in early and late replicating regions. Analysis of somatic mutations in early and late replicating regions showed that the higher mutation rate in late replicating regions disappears in tumors with defects in either mismatch repair (MMR) or global genome nucleotide excision repair (GG-NER) mechanisms^12,13^. These results suggest that the higher mutation rates in late replicating regions are due to less efficient repair in those regions. Interestingly, analysis of the association between different mutational signatures and replication properties revealed that most of the detected mutational signatures are significantly correlated with the timing or direction of DNA replication^8^, suggesting that the association between replication and mutagenesis is broad and involves many cellular mechanisms. Moreover, analysis of the association of RT with mutational signatures both in breast cancer^14^ and in multiple cancer samples^8^, revealed that some mutational processes are mainly associated with late RT.

It remains to be determined which mechanisms in late replication cause a higher load of mutations. Is the increased mutational load related directly to replication or is it a consequence of packaging in closed/less accessible chromatin? A recent paper addressed this issue and suggested that the association between mutation rates and RT is actually driven by the association between mutation rates and chromatin accessibility^15^. In contrast, it is clear that the replication process itself contributes to mutation rates, since there are several signatures that are associated with either replication rate^7^ or with replication fork direction^8^, two replication features that are not directly associated with chromatin structure.

Here we readdressed the contribution of RT to mutational distribution by performing a systematic analysis of the association between RT and mutational signatures using 2,787 Whole Genome Sequenced (WGS) tumors which are available from the PCAWG^16^. Our analysis is more comprehensive than previous analyses in several aspects. First, it expands previous analyses to the newest version of COSMIC SBS signatures (v3)^5^ which includes newly discovered signatures. Second, by developing a new context normalization method, we were able to expand our analysis to doublet base substitution (DBS) and small insertions and deletions (indels) signatures, which their RT association has not yet evaluated. Third, it analyzes both pan cancer and tissue specific signatures. Fourth, it distinguishes between the contribution of RT and of chromatin accessibility to mutagenesis. Taken together, our novel and comprehensive approach revealed that the association of late replication with higher mutation rate was an oversimplification, since many mutational processes are more frequent in early RT. This realization is important for better understanding of the etiology of many mutational signatures, which may lead to better prevention, diagnosis and treatment of cancer^17–19^. Indeed, improving signature identification by focusing on genomic regions in which each signature is more dominant, enabled us to better identify liver tumors harboring SBS16, and to reveal that it is a signature with potential clinical implications.

## Results

### Mutational profiles of tumors are different between early and late replication timing regions

In order to explore the relation between mutational processes and replication timing (RT), we compared mutation types in early vs. late replicating portions of the genome. To avoid variations of RT between tissues we restricted our analyses to the constitutive RT portions of the genome, which have a similar RT across a panel of 26 distinct human tissues^20^, and are similar also in cancer cells (**Supplementary Figure 1**). In total, 706Mb and 583Mb were analyzed as constitutive early and late regions, respectively (**Methods**). Somatic single nucleotide variants (SNV) of 2,787 WGS samples from the PCAWG project were analyzed for mutational signatures, separately for constitutive early replicating regions (ERR) and constitutive late replicating regions (LRR). Trinucleotide counts of each region were normalized relative to the trinucleotide distribution of the entire genome to account for differences in the trinucleotide distribution of ERR and LRR. Pan cancer analysis of 66 single base substitution (SBS) signatures across all tumors allowed testing for association of each signature with RT. The test is based on the mean difference (delta) between relative contribution of each SBS signature in ERR and LRR, across tumor samples, corrected for trinucleotide distribution (**Methods**; **Figure 1a)**. As expected^8,11^, we found many mutational signatures enriched in LRR (e.g. SBS signatures 1, 4, 8). Surprisingly, we also found many signatures enriched in ERR (e.g. SBSs 5, 16, 40). We repeated the analysis for each cancer project separately (see examples in **Figure 1b**). In almost all cases the RT bias was similar across cancer types (in which the signature exists), suggesting that the difference in SBS exposure in different RT regions does not depend on cancer type (**Figure 1c)**. SBS signatures such as SBS8, and to a lesser extent SBSs 2+13 (APOBEC-related), 5, 39, 40 and several others, demonstrated RT association in numerous cancer types. Furthermore, the cancer-type specific analyses emphasize the association between RT and mutational signatures of many tumor type specific signatures: SBS7a+b in melanoma, which are correlated with LRR and ERR respectively; SBS9 with LRR in lymphomas and Chronic Lymphocytic Leukemia (CLL); and SBS17a+b with LRR in gastric and esophageal cancer (**Figure 1b-c**).

**Figure 1.**
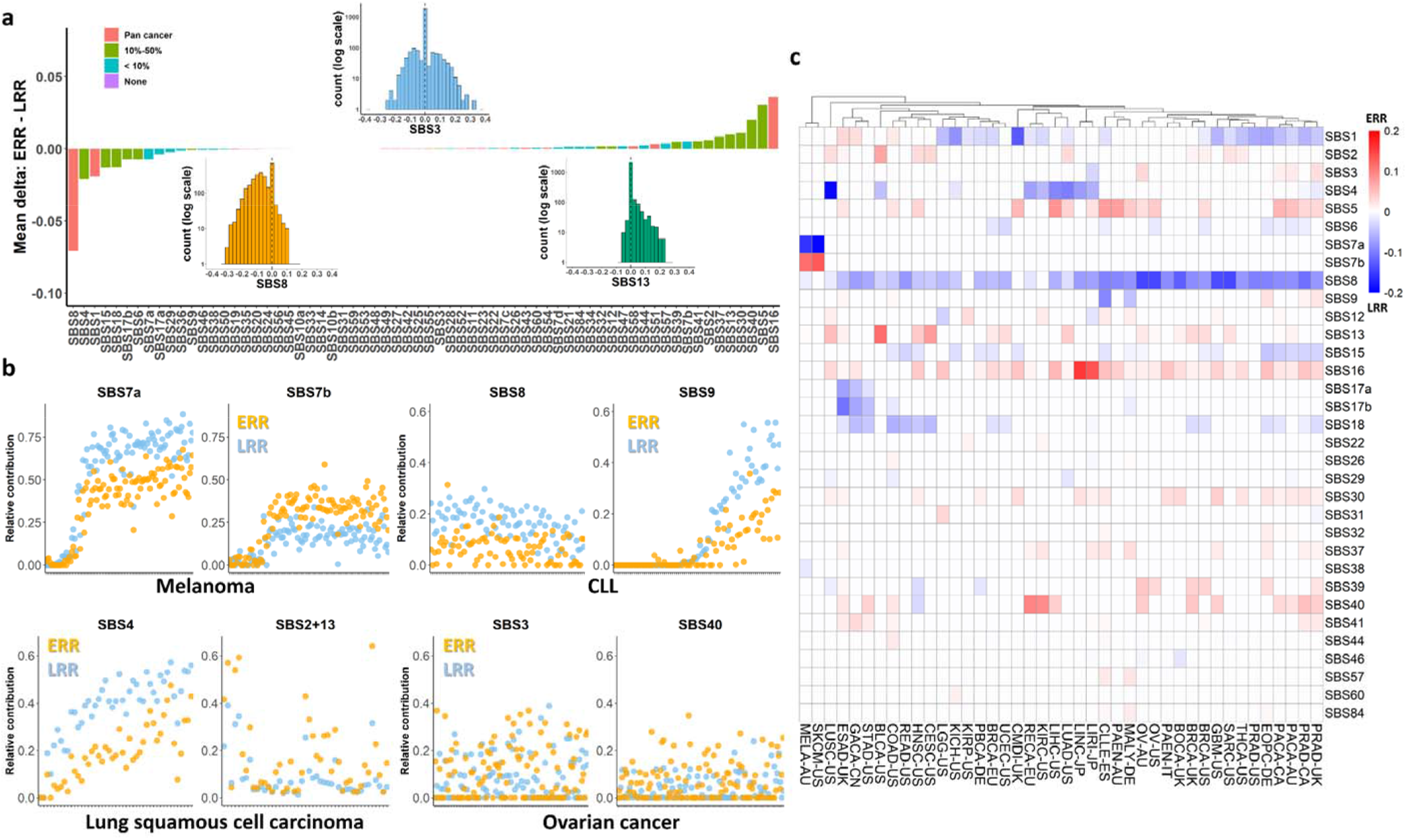
Replication timing contribution to single base substitution mutational signatures. **a,** Average replication-timing biases across all SBS signatures in 2,519 tumors. Shown is the delta between ERR and LRR. Positive value implies higher contribution in ERR, negative value implies higher contribution in LRR. Color of bars indicate in how many cancer projects the bias was found statistically significant (p < 0.05, FDR corrected Wilcoxon rank sum test). Signatures with RT bias in more than 50% of cancer-projects were considered as pan-cancer. Small histograms illustrate the distribution of the delta for three signatures. **b,** Scatter plots showing the contribution of the indicated signatures in ERR (orange) and LRR (blue) for individual cancer samples (X axis) in different cancer projects: melanoma MELA-AU and SKCM-US (SBS7a and SBS7b, left upper), CLL CLLE-ES (SBS8 and SBS9, right upper), lung LUSC-US (SBS4 and SBS2+13, left lower) and ovarian cancer OV-AU and OV-US (SBS3 and SBS40, right lower). X axis are tumors. **c,** Heatmap showing each SBS signature’s RT bias (p < 0.05, FDR corrected Wilcoxon rank sum test), or no bias (white), stratified by cancer types. Red and blue indicate positive and negative delta respectively. The projects were ordered using hierarchical clustering. Signatures with no bias in any project are not shown.

### Artificial genomes-based method for sequence-context correction

Since early and late replicating regions differ in their trinucleotide compositions and GC content^10^, differences in mutation distributions may stem from differences in the normal sequence context in those regions. Normalizing RT regions by trinucleotide counts compared to that of the genome counts addresses this issue, however it is difficult to apply it for more complex nucleotide context mutational events, as with indels. To this end, we developed an alternative normalization method which is based on the creation of an artificial sample for each tumor sample, in which the nucleotide context of each mutation is maintained but its genomic position is randomly chosen (**Methods**). These artificial genomes can be used for distinguishing between sequence context (which is maintained) and other biological processes that affect mutations distribution (that are active only in the original tumors). To validate this method, we compared the context-controlled RT bias, calculated by subtracting the bias in the artificial genomes from the bias in the un-normalized tumors, with the bias calculated using the trinucleotide normalization method. Overall, the results are highly similar (R = 0.960, p < 10^-16^, Pearson’s correlation test; **Figure 2a-b, Supplementary Figure 2).** This was true also when comparing the two normalization methodologies in the cancer-type specific analyses. The mean Euclidean distance between the ERR-LRR deltas of signatures between the two methods within the same project is 0.049, with an interquartile range of 0.040, significantly lower than the distances between different projects (mean distance of 0.145, p < 10^-16^, two-sided Student’s t-test; **Figure 2c)**.

**Figure 2.**
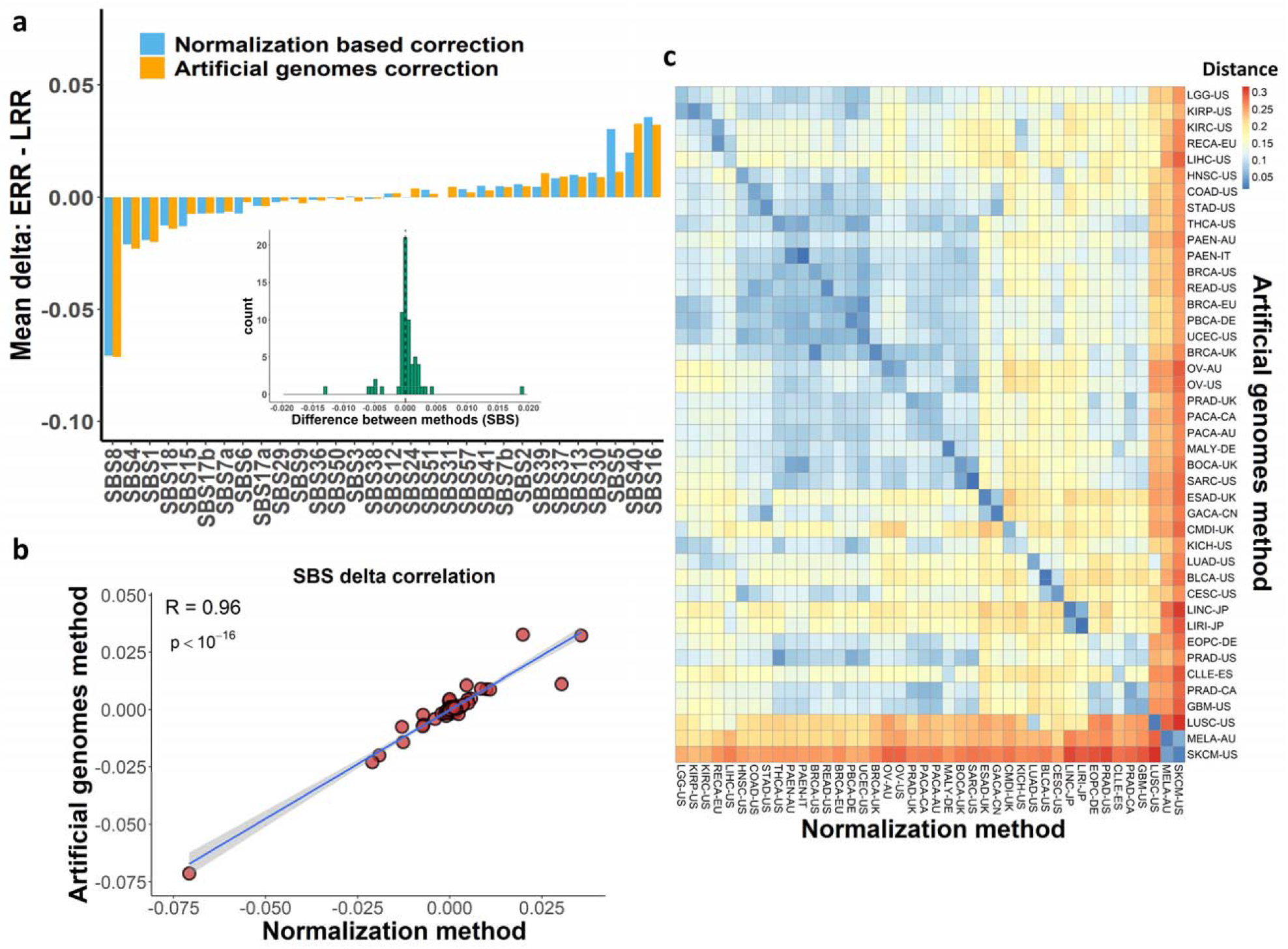
Artificial genomes-based method for sequence-context correction. **a,** Bar plot showing average replication-timing biases in normalization-based correction (blue) compared to the artificial genomes-based correction (orange) methods in SBS signatures across 2,519 tumors. Positive value implies higher contribution in ERR, negative value implies higher contribution in LRR. Heatmaps comparing all signatures, including many with delta of ~0, are available in the supplementary information (**Supplementary Figure 2b**). Small histogram illustrates the distribution of the differences between the deltas of the 2 methods. **b,** Scatter plot showing the correlation (R = 0.96, p < 10^-16^, Pearson’s correlation) between the deltas in the two methods. Each point marks SBS signature’s deltas. Regression line was calculated using ‘lm’ method in ggplot2. The grey shade area represents the 95% confidence interval. **c,** Euclidean distances heatmap between deltas’ distribution in cancer projects in the different methods. The diagonal represents the distances between same project in two methods. Lower values mean greater similarity. Projects are ordered by hierarchical clustering of the artificial-genomes method deltas. The deltas of each method separately are shown in **Supplementary Figure 2b.**

### Doublet base substitution and indels signatures

In addition to SBS there are two other types of small mutations – doublet base substitutions (DBS) and small insertions and deletions (indels) for which signatures were recently determined. DBS are less common events than SBS and indels, but were shown to be indicative of commonly occurring, single mutagenic events^5^. Our artificial genomes-based methodology for sequence correction allowed expanding our research to DBS and indels, whose association with RT has not been evaluated comprehensively thus far.

DBS signatures are based on the classification of 78 possible doublet-based mutation types. Overall, 11 DBS signatures were introduced by COSMIC, of which 8 were first reported recently^5^. Because of their rarity, only 163 tumor samples from ICGC and 139 samples from TCGA met our inclusion criteria for the RT analysis (i.e. at least 20 mutations in ERR and in LRR, and 90% success of reconstruction by the DBS signatures - see **Methods**). These 302 samples were mainly from three cancer types (83 melanomas; 123 liver cancers and 67 lung cancers). We applied our signatures analysis approach using the artificial genomes correction for ERR and LRR and found three signatures with RT bias (**Figure 3a**). DBS1 and DBS4 were enriched in ERR whereas DBS2 was enriched in LRR. The project specific analyses confirmed pan-cancer results and added two signatures enriched in ERR – DBS7 and DBS9 in liver and lung cancer, respectively. Finally, DBS11 showed surprising results since it is enriched in LRR in melanoma samples and in ERR in lung squamous cell carcinoma samples (**Figure 3a-b**).

**Figure 3.**
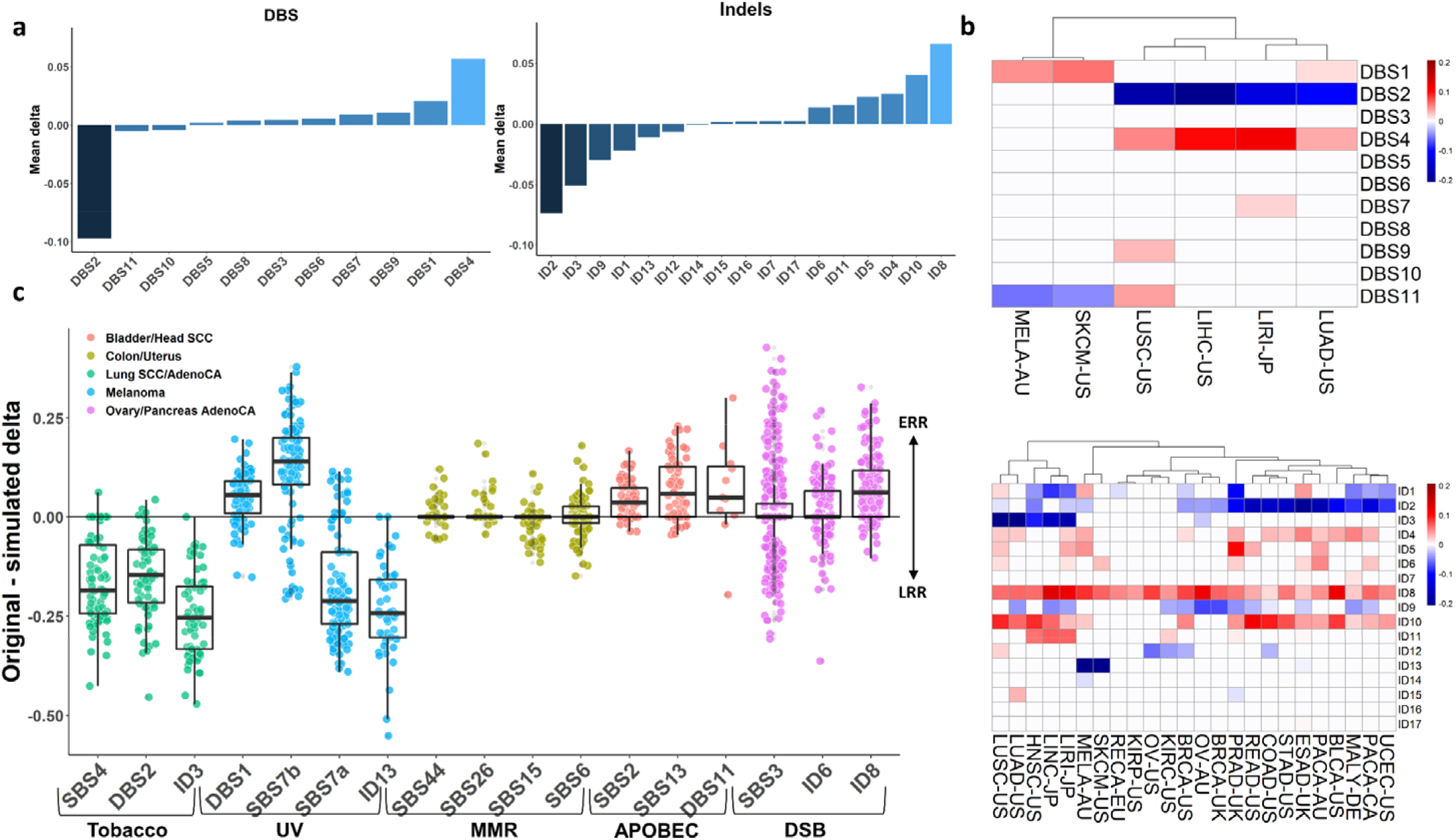
Replication timing contribution to doublet base substitution and indels mutational signatures. **a,** Bar graphs showing the RT bias (tumor RT delta – artificial genome RT delta) in relative contribution of relevant signatures for DBS (left) and indels (right) signatures. **b,** Heatmap delineating RT biases by different cancer types. Positive (Red) and negative (blue) values indicate ERR bias and LRR bias, respectively. All colored cells have p < 0.05, FDR corrected Wilcoxon rank sum test. Upper: DBS signatures; Lower: indels signatures. **c,** Distribution of the contribution of RT (tumor RT delta – artificial genome RT delta) for selected SBS, DBS and indels signatures clustered by etiology. Each point marks the difference between tumor delta and artificial genome delta of a given sample. Points are colored by cancer type and a specific etiology. Cancer projects presented are those in which we anticipate to find the selected signatures^5^. In boxplots: the center line marks the median value, upper and lower limits mark first and third quartiles and the whiskers cover data within 1.5× the IQR from the box.

The classification of small insertions and deletions signatures (ID signatures) is based on 83 subtypes of indels events. In total, we analyzed 891 samples that met the inclusion criteria (i.e. at least 100 indels in ERR and in LRR, and 90% success of reconstruction by the ID signatures - see **Methods**). Pan-cancer analysis revealed a few signatures with strong RT biases, including ID2, ID3, ID9 and ID1 in LRR; and ID8, ID10, ID4 and ID5 in ERR (**Figure 3a-b**). Cancer type specific analyses confirmed these results and revealed the association of ID13 with LRR in skin cancers (**Figure 3b**).

These results are consistent with the single substitutions results **(Figure 3c)**. SBS4, DBS2 and ID3 are associated with tobacco damage and found enriched in LRR. Similarly, APOBEC related signatures (SBS2, SBS13 and DBS11) are associated with ERR, as have been recently suggested^21^. Both DBS1 and ID13 are associated with UV damage but they differ in their enrichments. While DBS1 resembles SBS7b in its enrichment in ERR, ID13 resembles SBS7a in its association with LRR. This can be explained by the type of pyrimidine dimer characteristic of each signature. In ID13 and SBS7a the first residue in the dimer is T (TT and TC, respectively) and the bias is toward LRR, whereas in DBS1 and SBS7b the first residue is C (CC and CT, respectively) and the bias is toward ERR (**Figure 3c**). Further studies are needed to understand the repair mechanisms that are responsible for these differences.

We found that all three DNA double strand break (DSB) associated signatures (SBS3, ID6 and ID8) are enriched in ERR (**Figures 1a, 3a-c**). This finding goes along with the higher prevalence of chromosomal rearrangements (which are the consequence of unrepaired DSBs) in ERR (reviewed in^11^), and by the analysis of End-seq data^22^, which revealed significantly more DSBs in ERR than in LRR (**Supplementary Figure 3**).

Taken together, the consistent association between replication timing bias of related mutational signatures (SBS, DBS and indels) and the variable association between the processes, further support the realization that different mutational processes vary in their association with RT.

### The tangled relationship between RT and chromatin activity

We have shown a clear association between RT and many mutational signatures, which goes beyond the differences in sequence distribution between ERR and LRR. RT is associated with chromatin structure - ERR are usually more accessible and active than LRR^10^. Thus, it is possible that RT is just a proxy for chromatin structure, which affects mutation by modulating DNA accessibility and the RT itself may not contribute to mutational processes. Indeed, a recent publication suggests that mutation rates and distributions are mainly associated with chromatin structure and less with RT^15^. In order to directly address this question, we wanted to define four relatively uniform genomics regions: ERR-active, ERR-inactive, LRR-active and LRR-inactive and compare the contribution of mutational signatures in each of those regions. To this end, we took advantage of a recent annotation that used an entropy-based approach to assign a chromatin state to each genomic region across all cell types profiled in the Roadmap Epigenome Consortium^15^. In order to get sufficient genome coverage, we profiled the RT of several cancer cell lines (**Methods**). Together, the profiled cancer cell lines cover most of the signatures with a clear RT bias (|delta|>0.05 in at least one project; **Figure 1b**). Intersecting active and inactive domains with RT data allows us to create the 4 genomic categories **(Figure 4a; Supplementary Table 2; Methods**). Using these categories, we repeated our signatures analyses.

**Figure 4.**
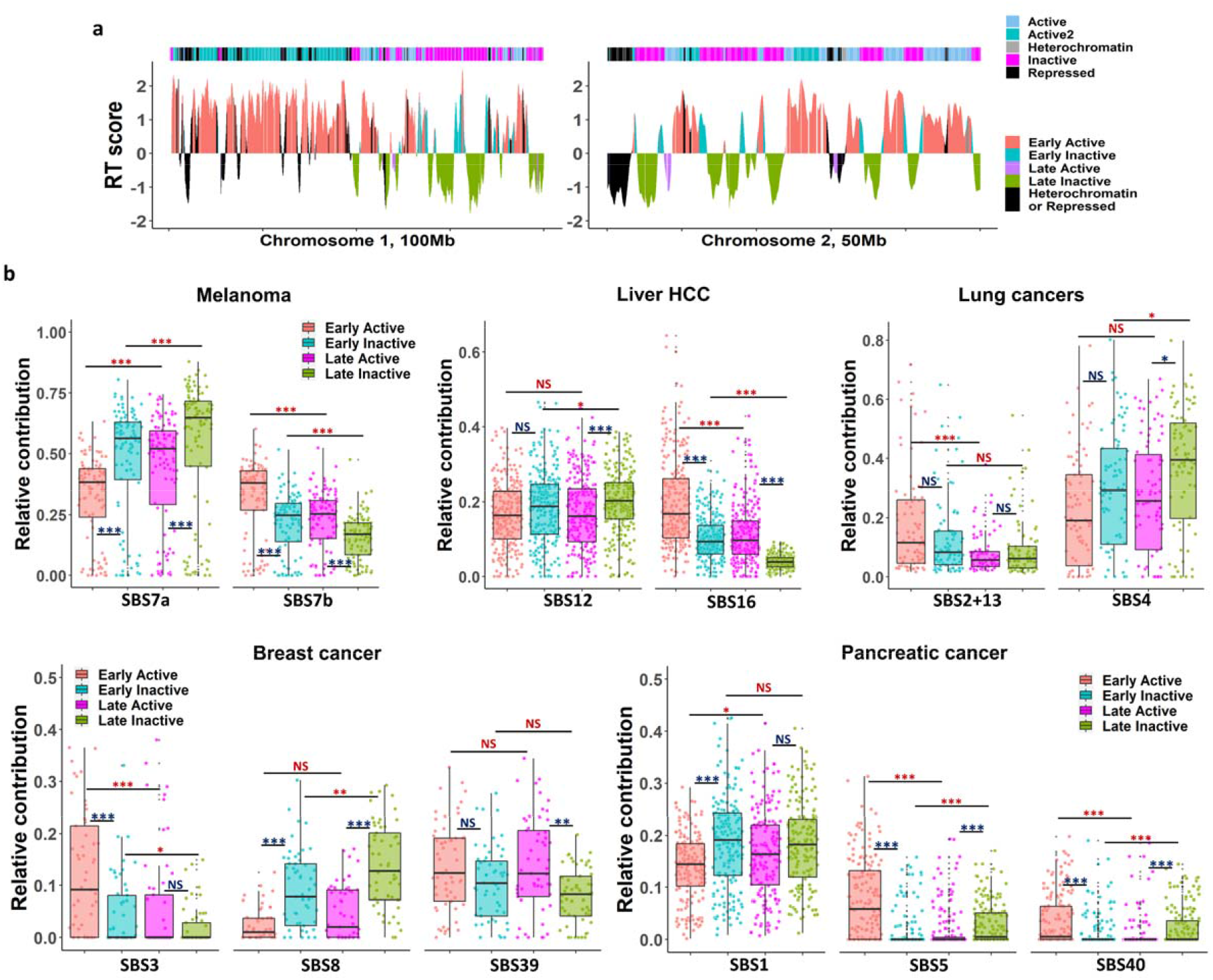
Impact of replication timing and chromatin activity on different mutational processes. **a,** RT and chromatin activity regions distributed across 100Mb in chromosome 1 (left) and 50Mb in chromosome 2 (right) in a melanoma cell line. Upper bar illustrates chromatin activity annotations taken from^15^. Y-axis represents the replication timing score of each area. The colors capture the four categories used for the analysis and regions excluded from the analysis (black). **b,** Relative contribution of selected signatures, stratified by RT and chromatin regions. All p-values derived from FDR corrected Wilcoxon’s rank sum test. NS, not significant; *, p < 0.05; **, P < 0.01; ***, p < 0.001. Red and dark-blue font colors indicate RT regions comparisons (i.e., ERR vs LRR) and chromatin accessibility comparisons (i.e., active vs inactive) respectively. In boxplots: the center line marks the median value, upper and lower limits mark first and third quartiles and the whiskers cover data within 1.5× the IQR from the box. Each dot represents a single tumor. Left upper: SBS7a/b in melanoma samples (n=108); Middle upper: SBSs 12 and 16 in liver hepatocellular carcinoma (n=257); Right upper: SBS2+13 (APOBEC related) and SBS4 in lung adenocarcinoma and squamous cell carcinoma (n=84); Left lower: SBSs 3, 8 and 39 in breast invasive carcinoma (n=57); Right lower: SBSs 1, 5 and 40 in pancreatic carcinoma (n=176).

We found that RT contributes to the creation of UV-related mutational signatures independently of chromatin activity, as both SBS7a and SBS7b showed significantly different contribution in LRR and ERR, even when the domain activity was controlled for (all p-values < 10^-4^, FDR corrected Wilcoxon’s rank sum test; **Figure 4b).** Interestingly, the opposite was true as well – domain type contributes to UV related mutagenesis even when RT was controlled for (all p-values < 10^-5^).

Overall, this phenomenon was repeated in various signatures. SBSs 16, 2+13, 3, 5 and 40 showed highest enrichment in early-active regions in liver, lung, breast and pancreatic cancers respectively, statistically significant more than when one factor was controlled, i.e. in late-active or early-inactive regions **(Figure 4b).** Similarly, SBS8 and to a lesser extent also SBS4 showed highest enrichment in late-inactive regions in lung and breast cancer, respectively, statistically significant more than when one factor is controlled **(Figure 4b).** In contrast, other signatures show higher dependency on chromatin accessibility than on RT. This is true for example for SBS39 in breast cancer that its association with ERR **(Figure 1)** is actually association with accessible chromatin **(Figure 4b)**. Similarly, the association of SBS1 to LRR in pancreatic cancer can be mainly explained by association with the inactive parts of the genome **(Figure 4b)**. Taken together, our results clearly demonstrate that both RT and chromatin accessibility contribute independently to mutagenesis with different effects on different mutational processes.

### Focused signatures analysis approach reveals association of SBS16 and survival rates

Finding the uneven genomic distribution of many signatures (summarized in **Supplementary Table 1**), suggests that taking RT information into account may improve mutational signatures identification. We therefore developed a focused signature analysis approach, and demonstrated its power by studying the contribution of SBS16 to liver cancer.

SBS6 is a signature of unknown etiology, found mainly in liver cancer^5^. It was shown to be associated with male gender, alcohol use and tobacco consumption^23^. Here, we showed that SBS16 is highly enriched in early replicating regions (**Figures 1a, 1c, 5a**). In light of our findings, we explored the contribution of SBS16 in liver cancer, using either the entire genome or focusing on ERR only. Out of 257 samples in the LIRI-JP project we found only 58 samples with a significant (>10%) relative contribution of SBS16 using the entire genome for signature identification, while almost three times more samples (170 samples) were found with SBS16 signature analysis focused only on ERR (**Figures 5b-c**). Since our approach nearly tripled the number of tumors harboring SBS16, we further characterized those tumors and designated three groups: i) SBS16 positive in both analyses (N=58); ii) SBS16 positive only in ERR-focused analysis (N=122); and iii) SBS16 negative (less than 10% contribution) (N=69). Gender analysis of the three groups revealed that the two SBS16 positive groups are mainly composed of males (53/58 in whole genome group, 99/122 in ERR only group), a statistically significant indication that both groups are indeed SBS16 positive (in a cohort of 190/257 = 74% males, the likelihood that randomly chosen 122 samples will contain 99 males is p < 0.05, one-sided exact binomial test). In contrast, in the SBS16 negative group there are 52% females and 48% males.

**Figure 5.**
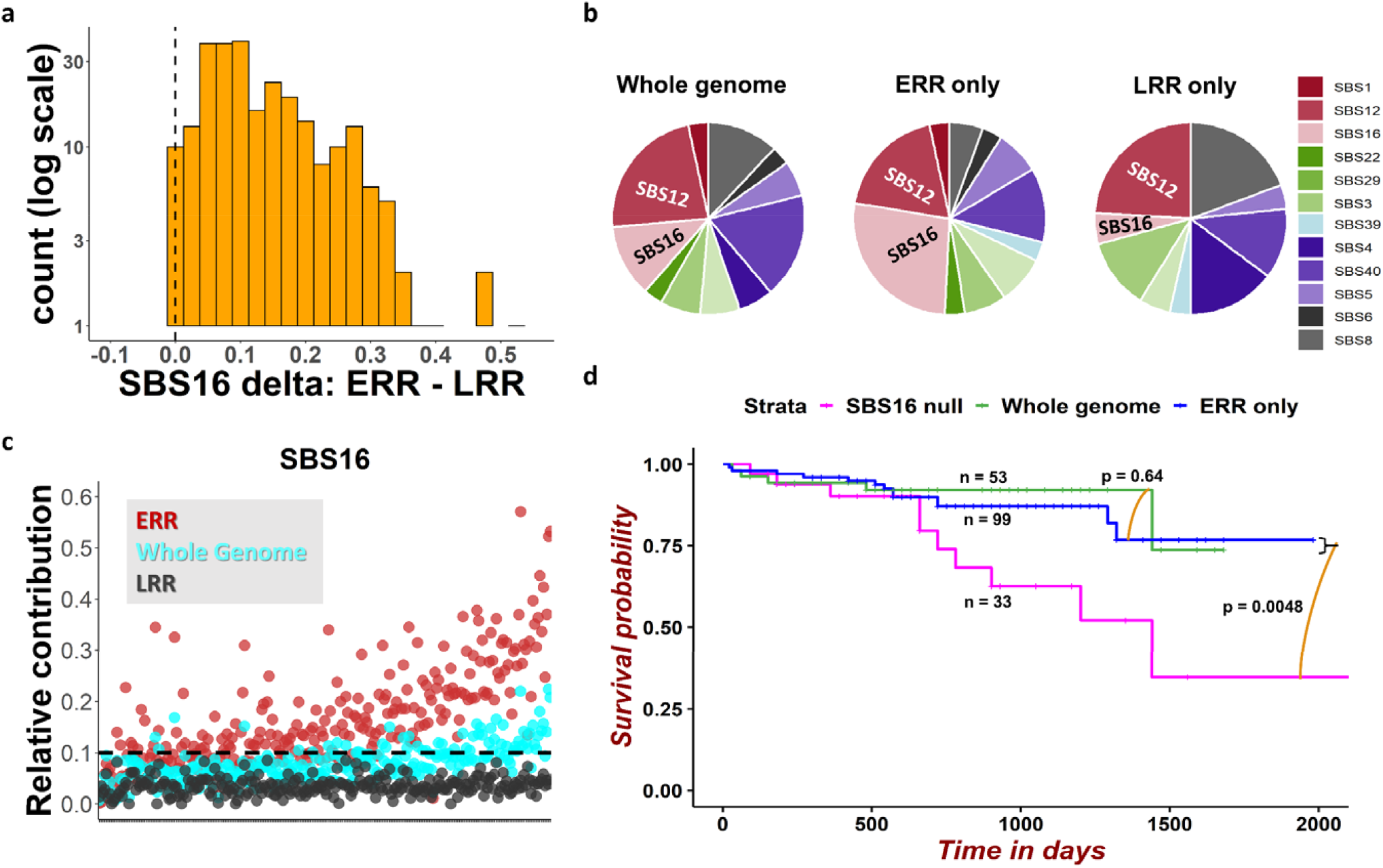
Better survival of liver patients harboring SBS16 revealed by focused signatures analysis identification. **a,** Histogram of SBS16 delta (ERR – LRR contribution). Positive values indicate higher contribution in ERR. **b,** Pie charts illustrating the changes of SBS signatures contribution in entire genome, ERR only, and LRR only. The average contribution of each signature in the LIRI-JP project is presented. **c,** Scatter plots showing the contribution of the SBS16 in ERR (red), whole genome (cyan) and LRR (black) in individual cancer samples. X axis are tumors. **d,** Survival analysis for three groups of males with liver hepatocellular-carcinoma from LIRI-JP project: green – harboring the SBS16 signature in entire genome analysis (n=53); blue - harboring the SBS16 signature only in ERR analysis (n=99); magenta – does not contain the SBS16 signature (n=33). P-values demarcate SBS16 in ERR only vs. SBS16 in whole genome (upper, p=0.64) and combined SBS16 positive group (ERR only + whole genome, n=152) vs. no SBS16 group (lower, p=0.0048). P-values derived from Cox proportional-hazards regression model.

Survival analysis of male patients revealed that both SBS16 positive groups have similar survival times, which are significantly longer compared to the survival rates of the SBS16 negative group (P = 0.64 and P = 0.0048, respectively, Cox proportional-hazards regression model; **Figure 5d**). This difference in survival cannot be identified using the classical entire genome approach, since 99 tumors, which are SBS16 positive only in ERR-focused approach and thus have better survival rates, are considered SBS16 negative while exploring all genome **(Supplementary Figure 4a; p = 0.25).** Males survive longer than females in LIRI-JP project (**Supplementary Figure 4b**). However, we observed that the better survival is restricted only to individuals harboring SBS16, whereas for individuals without SBS16 the survival was similar between males and females (P=0.66, Cox proportional-hazards regression model; **Supplementary Figure 4b**). Taken together, focusing signature analysis to the relevant parts of the genome improved the identification of SBS16 and discovered its association with better survival. These results were not a consequence of the threshold we chose (10% contribution), since similar results were obtained for a large range of thresholds (**Supplementary Figure 5**).

As expected, applying the focused signature analysis approach to many other signatures with clear association with RT increased the number of tumors harboring each signature **(Supplementary Figure 6)**.

## Discussion

A great deal of evidence shows higher mutation rates in late replicating regions^11^, however the mutational processes that lead to this bias are not yet clear. By using mutational signature analyses separately on genomic regions that replicate at either early or late S phase we were able to identify many mutational processes that are differentially associated with RT **(Supplementary Table 1)**.

The association between RT and mutational processes has been previously addressed – perturbation either to the mismatch repair (MMR) or to the global genome nucleotide excision repair (GG-NER) mechanisms abolishes the differences between LRR and ERR, suggesting that both these repair processes are more efficient in ERRs^12,13^. This specific mechanism probably stems from the fact that ERR genomic regions are packed in open chromatin that allows better accessibility to the repair proteins. More recently, a systematic analysis of mutations distribution revealed that many mutational signatures are associated with RT, suggesting that additional processes (beside MMR and GG-NER) show differential efficiencies in different genomic locations^8^. This comprehensive study found that most of the signatures that show association with RT were enriched in LRR, and only two signatures - SBS5 and SBS16, were enriched in ERR. Recently it was shown that another mutational process - APOBEC3 mutagenesis, is enriched in ERR^21^.

These findings encouraged us to readdress the possibility that certain mutational processes are enriched in the ERRs and that the association between mutation rate and RT is more complex than just having a greater mutation load in the late replicating and closed portions of the genome. We improved previous analyses in several ways. First, we restricted our analyses to genomic regions in which the replication timing was found to be constitutive^20^, minimizing the effect of variation in RT between tumor types. Second, we used relative contribution in order to control for different mutation rates in different tumors and tumor types. Third, we used the latest version of COSMIC mutational signatures (version 3), which is much more precise then previous versions regarding contamination between different signatures and increased the number of known SBS signatures substantially^5^. Fourth, we expanded the analysis to DBSs and indels related signatures. Fifth, we performed pan-cancer analysis as well as cancer type specific analysis. This identified pan cancer mutational processes as well as cancer specific processes and thus enabled identification of the RT association of rare signatures. Finally, we corrected for differential sequence distributions in ERR and LRR both by normalizing for trinucleotide frequencies and by developing an artificial genomes-based approach, which is crucial for considering differential nucleotides sequence distribution for DBS and indels.

Taken together our results revealed the RT association of each mutational signature. We found that in contrast to the previous perception of general preferences for mutagenesis in late replicating regions^7,8,11,12^, the actual picture is more subtle with different mutational processes showing preferences to either ERR, LRR or no association with RT. The most obvious example of this new perception is the difference between SBS7a and SBS7b. Both signatures are caused by UV damage but differ in their actual signatures most probably due to differences in yet unknown repair mechanisms, still 7a is enriched in LRR whereas 7b in ERR. The generation of artificial genomes allowed us to delineate context-bias of signatures, as well as to determine the RT bias of DBS and indels **(Figure 2)**. We demonstrated that knowledge of the mutation distribution bias across genomic regions can improve the sensitivity of signature analyses by focusing on the more relevant parts of the genome. This realization allowed us to better identify liver tumors with SBS16 and to show for the first time a strong association between SBS16 and improved survival rates.

What causes the non-uniform distribution of mutational signatures along the genome? One possibility is that mutations are derived by sequence distribution (for example one expects to find more APOBEC related signature SBS13 in genomic regions rich with TCT or TCA trinucleotides). The other possibility is that the distribution of either the damage or the repair processes is not uniform and certain genomic regions are more susceptible to these specific processes. For example, as has been shown before, both MMR and NER are more efficient in ERR^12,13,21^. Our analyses support all three possibilities – some mutational signatures show association with RT mainly due to sequence distribution (SBS3 and 12, **Supplementary Figure 2a**). Other cases, such as the UV response signatures 7a and 7b show opposite association with RT, in spite of relative uniform distribution of UV damage^24^, suggesting that they differ in the cellular response to the damaged DNA. SBS7a (and ID13 as well) shows higher contribution in LRR, whereas SBS7b and DBS1 are enriched in ERR. These different preferences are probably attributed to different repair mechanisms active in ERR and LRR. Indeed, trans-lesion synthesis polymerases show differential activity throughout the cell cycle^25^, raising the possibility that different TLS polymerases are active during early and late S. Finally, the higher association of both ID6 and ID8 to ERR is probably due to higher frequency of DSBs in ERR, which was directly observed by the End-seq methodology that provides a landscape of DNA double-strand breaks prior to DNA repair (**Supplementary Figure 3)**.

RT is strongly associated with chromatin structure - ERR are gene rich, arranged in an open chromatin and have higher transcription rates, whereas LRR are enriched within the closed and silenced portions of the genome^10^. Thus, our findings of the association between RT and mutational signatures do not necessarily imply a direct association between replication and mutagenesis and it may be actually an association between the chromatin structure and the mutagenesis processes. Indeed, a recent paper suggests that the association between mutation rates and RT is actually an association between mutation rates and chromatin accessibility^15^. On the other hand, the contribution of the replication process itself to mutation accumulation is clear since several signatures are directly associated with replication rate^7^ or with replication fork direction^8^. These two options are not mutually exclusive and there are probably some signatures that are affected by accessibility whereas other signatures are associated with actual replication processes that are different between early and late S phase. We directly addressed this issue by dividing the genome into four regions with relatively uniform RT and chromatin accessibility. This stratification allowed us to test separately the effects of RT and chromatin accessibility on mutation rates for many mutational signatures. We found that for many signatures (such as SBSs 3, 2+13, 7a, 7b, 16, 5, 40, 8 and 4) both chromatin accessibility and RT contribute separately to the mutational activities. On the other hand, in other signatures (such as SBSs 1 and 39) the contribution of chromatin accessibility was much higher (**Figure 4**). Our results suggest that the interplay between RT and chromatin structure is tangled, and one genomic feature cannot fully explain the mutational landscape of the other. We showed that RT contributes to mutational rates of different mutational processes independently of chromatin accessibility, and vice versa.

Our findings have possible clinical implications. Focusing on specific genomic regions based on RT can lead to higher sensitivity to detect mutational signatures. As an example, we have shown that analysis focused on ERR more than tripled the number of samples identified with SBS16. Importantly – the patients with SBS16 detected only in an ERR-focused analysis had better survival, which is comparable to the survival of patients with SBS16 detected in whole genome analysis. This shows that not only more patients were detected with SBS16 signatures but that the clinical implication of ERR focused SBS16 detection is similar to whole genome identification of SBS16.

Taken together, the present study shows that replication timing is associated with many mutational signatures and that at least some of this association is specific to RT related processes. We showed clinical implication of these findings. We believe that as genomic data from more tumors is accumulated, more associations will be identified and that this might lead to better use of signature information to treat cancer patients.

## Methods

### Data sources

We downloaded somatic mutation calls (VCF files) from the Pan-Caner Analysis of Whole Genomes (PCAWG) consortium release of 2,787 whole-cancer genomes and their matching normal tissue across 38 tumor types^16^. The data consists of two sources: The International Cancer Genome Consortium (ICGC) - 1,902 samples, and The Cancer Genome Atlas (TCGA) - 885 samples. Each source utilized its standard variant call pipeline (Consensus calls for ICGC, and the Broad institute variant calling pipeline for TCGA). The somatic mutation profile of the two consortiums were very similar both in terms of 96 trinucleotide context (**Supplementary Figure 7a**), and in terms of mutational signatures (**Supplementary Figure 7b**). Accordingly, we combined the mutation calls data for all analyses.

### Replication timing regions and chromatin annotations

In order to minimize the effect of variation in RT between cell types, we used only the constitutive RT regions for most of our analyses, which constitute approximately 40% of the human genome that have the same RT in 26 tissues examined^20^ and are also similar in cancer (**Supplementary Figure 1)**. We used the median RT of the genome to separate between early and late replication **(Supplementary Figure 8)**. Among the constitutive RT regions 706Mb are defined as ERR and 583Mb as LRR. For the stratification of melanoma, liver HCC, breast cancer, lung cancer and pancreatic carcinoma samples by chromatin activity, we used RT data of the entire genome (and not only the constitutive regions) of a cell line derived from the relevant cancer type. These RT profiles were intersected with chromatin annotation from a recent publication^15^. Regions annotated as either “active” or “active2” were defined as active chromatin whereas the “inactive” regions were defined as inactive chromatin. The use of the full genome RT profiles for these analyses was crucial, since early-inactive and late-active groups are small (80Mb and 17Mb out of 706Mb and 583Mb of constitutive RT regions respectively). Replicating timing of four cell lines was produced by us: melanoma cell line FM-55-P, breast cancer cell line BT-549, lung cancer cell line A549 and pancreatic carcinoma cell line PANC-1. RT data of liver HCC cell line HepG2 was downloaded from https://www2.replicationdomain.com/database.php#. In total, 860-950Mb were classified as early-active, 200-433Mb as early-inactive, 113-197Mb as late-active and 865-1,024Mb as late-inactive **(Supplementary Table 2).**

### Mutational signatures analysis

To measure resemblance between different regions in the same sample, we used cosine similarity:

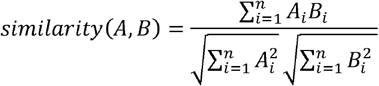

The cosine similarity values range between 0 (completely different) and 1 (identical).

Mutational profiles of SBS, DBS and indels were extracted using SigProfilerMatrixGenerator^26^. We excluded tumors with too few mutations (< 200 SBS, < 100 indels, < 20 DBS)^27^. Unless stated otherwise, for each region used in the paper aside for when using the method of comparison to artificial genomes, we normalized the mutation counts by the abundance of the trinucleotide context relative to that of the whole genome. SBS, DBS and indels signatures referred to in this article are all from the COSMIC database v3^28^. Overall, we used 66 SBS signatures, 11 DBS signatures and 17 indels signatures. Mutational signatures were detected for each sample and each region independently, by solving the Non-Negative Least Squares (NNLS) algorithm using MutationalPatterns R package^29^, fitting known COSMIC mutational signatures to the given mutational profile. This approach has been proposed for assessing signatures in well-studied cancer samples and when not aimed to discover new signatures as described^5,30,31^. The fitting approach may falsely assign too many signatures to a sample (signature “bleeding”). Thus, we used a-priori knowledge from the gold-standard COSMIC database for excluding non-relevant signatures as described in Alexandrov et al. and in Maura et al.^5,31^. Samples in which the similarity between the original mutational profile and the reconstructed mutational profile (given by the fitting algorithm results) were < 0.9 (cosine similarity) were excluded from further analyses. The contribution of each signature to the mutations in each tumor can be measured in absolute terms (counting the number of mutations attributed to each signature) or in relative terms (assessing the percentage of mutations attributed to each signature). We preferred the relative methodology, since it controls for different mutation rates in different tumors and tumor types.

### Artificial genomes

For each of the tumor samples, we generated 10 artificial genomes using SigProfilerSimulator^26^. The artificial genomes carry the exact same mutational profile (classified by 96 SBS subtypes, 78 DBS subtypes and 83 ID subtypes, including microhomology deletions) as the original sample (cosine similarity = 1). The simulated mutations are randomly distributed across the genome in an unbiased fashion. These artificial genomes were analyzed for signatures. The mean of the 10 ERR and LRR mutational signature profiles was used for the comparison with the samples’ actual signatures profiles.

### Delta analysis

The delta is the relative contribution of each signature in ERR minus its relative contribution in LRR. Relative contribution of each signature in each sample was calculated as number of mutations attributed for a specific signature in a specific sample, divided by the sum of mutations in that sample. Delta was calculated only if the contribution of a specific signature was more than 5% in at least one region (ERR or LRR). A signature was considered pan-cancer if it appeared significantly associated with RT (p < 0.05 for a specific signature in a specific cancer project, FDR corrected Wilcoxon rank-sum test), in at least 50% of projects included in the analysis. The delta analysis was performed in the same way for artificial genomes. Differences of deltas were calculated between real genomes and artificial genomes’ signatures deltas (i.e., signatures’ delta of the original genomes minus signatures’ delta of the artificial genomes). Only projects with at least 15 tumors, after applying the inclusion criteria (see above), were included in cancer type specific analyses.

### SBS16 detection and survival analysis for LIRI-JP project

Clinical data of patients from LIRI-JP project were obtained from the ICGC data portal. Since only few cases of female’s liver tumors contain SBS16, we restricted our survival analysis to males. Survival analysis was performed using R packages ‘Survival’ and ‘Survminer’. In Cosmic v3 SBS16 is a liver-specific well-defined signature^5^. Thus, one can reliably use a relatively low threshold for identifying SBS16 positive samples in liver cancer samples. We performed the survival analyses with a range of thresholds (8% to 12%). For each analysis, a sample was considered SBS16 positive/negative if SBS16’s relative contribution was above/below the specific threshold, respectively (**Supplementary Figure 5)**.

### Statistics

Statistical analyses were performed using R version 4.0.3. If not stated otherwise, the comparison of two distributions of continuous values was tested with a paired Wilcoxon rank sum test. For multiple comparisons, P-values were corrected by false discovery rate (FDR). All box plots are presented according to the standard box plot notation in R (ggplot2 package).

## Declarations

### Ethics approval and consent to participate

All of the participants were consented by previous profiling studies. This work utilized previously generated cancer-patients related datasets.

### Consent for publication

All authors have agreed to publish this manuscript.

### Availability of data and material

Sequence data reported in this study have been submitted to the Sequence Read Archive (SRA) under accession numbers PRJNA419407 and PRJNA726798. Other datasets utilized in this study are available online as described in the Methods section.

## Acknowledgments and funding

We thank Professor Ben Berman for assistance in data analysis. This work was supported by research grants from the Israel Academy of Sciences (grant # 184/16 to I.S., grant # 2479/20 S.R.), ISF-NSFC (grant #2555/16 to I.S.), the Israel Cancer Research Foundation (I.S.), the Binational Science Foundation (grant # 2019688 to I.S.), the joint fund for the Hebrew University and its affiliated hospitals (to I.S. and S.R.), Trudy Mandel Louis Charitable Trust (S.R.) and Hadassah-France (S.R.).

## Authors’ contributions

A.Y., I.S. and S.R. designed the study. A.Y. and O.V. performed the bioinformatic and computational analyses. O.V., A.G., D.J.M and A.K. performed the replication timing profiling of cancer cell lines. All authors discussed the results and commented on the manuscript.

