## Supplementary information for "Cancer mutational processes vary in their association with replication timing and chromatin accessibility"

### Supplementary Figure 1

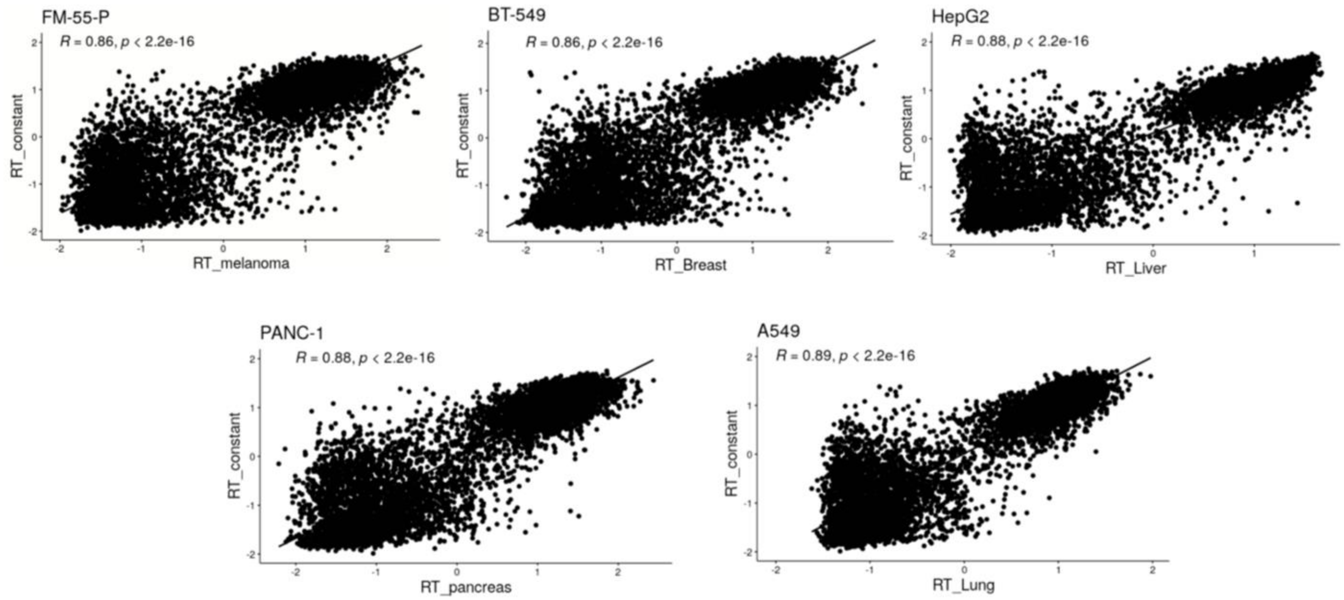


**Supplementary Figure 1 |** Correlation between replication timing (RT) in cancer cell lines (X axis) and constitutive RT regions (y axis). All R > 0.85, all p < $10^{-16}$, Pearson's correlation test. Upper: FM-55-P (melanoma, left), BT-549 (breast cancer, right) and HepG2 (liver cancer, right). Lower: PANC-1 (pancreatic carcinoma, left) and A549 (lung carcinoma, right).

### Supplementary Figure 2

**a**

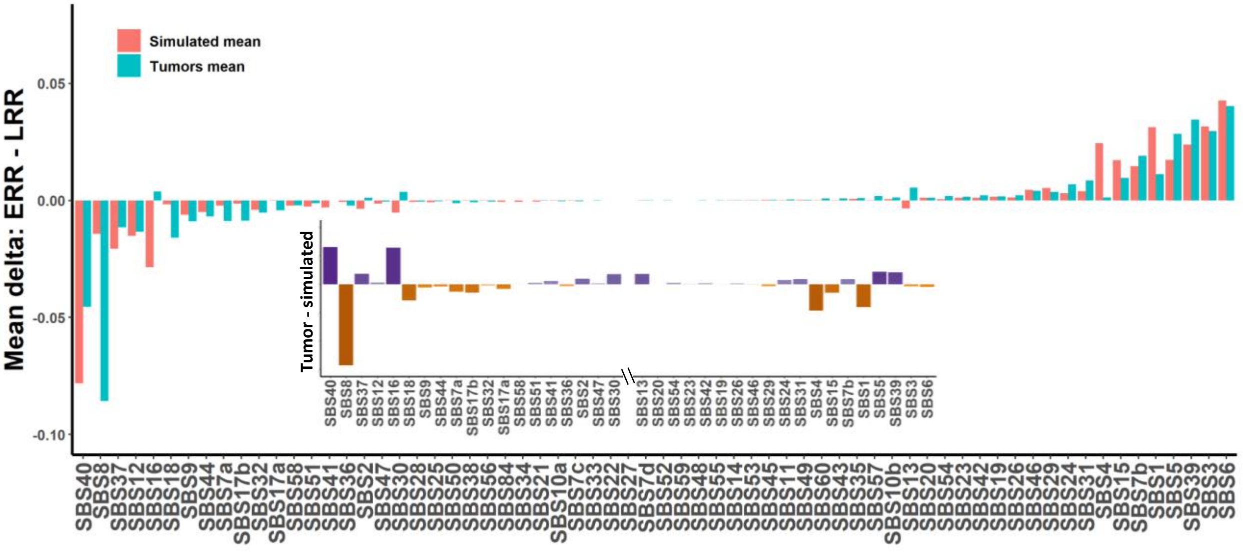

**b**

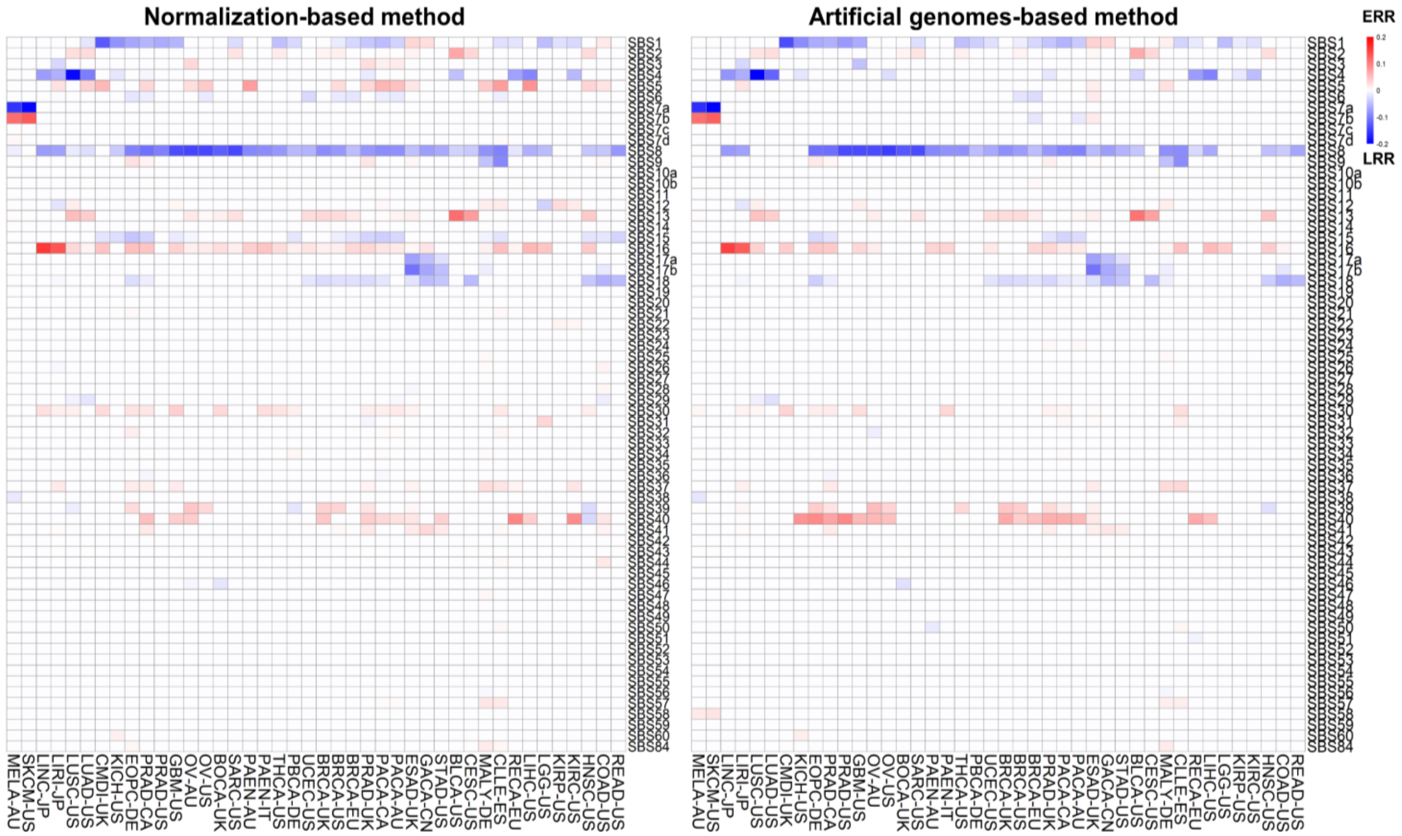


**Supplementary Figure 2 | a,** Bar plot showing average replication-timing biases in un-normalized tumors (blue) and in artificial genomes (red) in SBS signatures across 2,519 tumors. Positive value implies higher contribution in ERR, negative value implies higher contribution in LRR. The small bar plot illustrates the differences between the original and the artificial, i.e. the sequence-context independent RT bias. **b,** Heat maps of RT association of 66 SBS mutational signatures per cancer type. Red and blue values imply ERR and LRR bias respectively. All colorful values represent $p < 0.05$, FDR corrected Wilcoxon rank sum test. Left: Bias revealed by trinucleotide counts normalization relative to that of the entire genome. Right: Bias revealed by artificial genomes-based method, i.e. contribution differences (delta) in ERR and LRR in un-normalized genome against the delta in the artificial genomes. Note the similarity between the two heat maps.

### Supplementary Figure 3

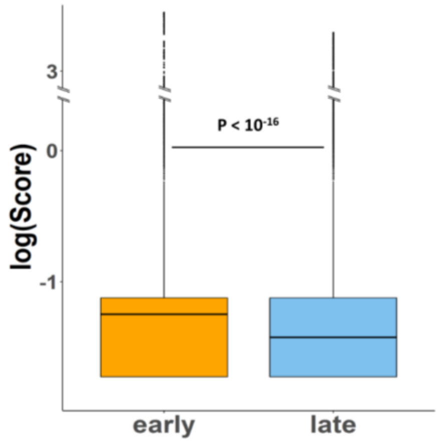


**Supplementary Figure 3 |** Boxplots showing the distribution of End-seq scores separately for End-seq peaks located at ERR or LRR. P-value derived from a Wilcoxon rank sum test.

### Supplementary Figure 4

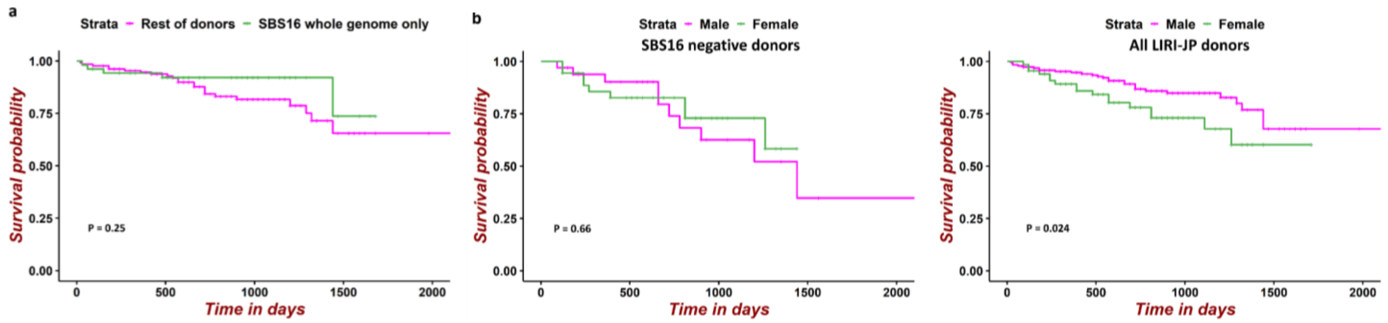


**Supplementary Figure 4 | a,** Survival analysis using entire genome approach in LIRI-JP project. SBS16 negative male patients (<10% contribution in the entire genome, n=132) vs. SBS16 positive samples (>10% contribution, n=53) in the entire genome. **b,** Survival analysis, SBS16 negative ( <10% contribution in ERR) male (n=33) vs. female (n=36) donors (left; p = 0.66, Cox proportional-hazards regression model) and survival analysis of male (n=190) vs. female (n=67) in entire LIRI-JP project (right; p = 0.024).

### Supplementary Figure 5

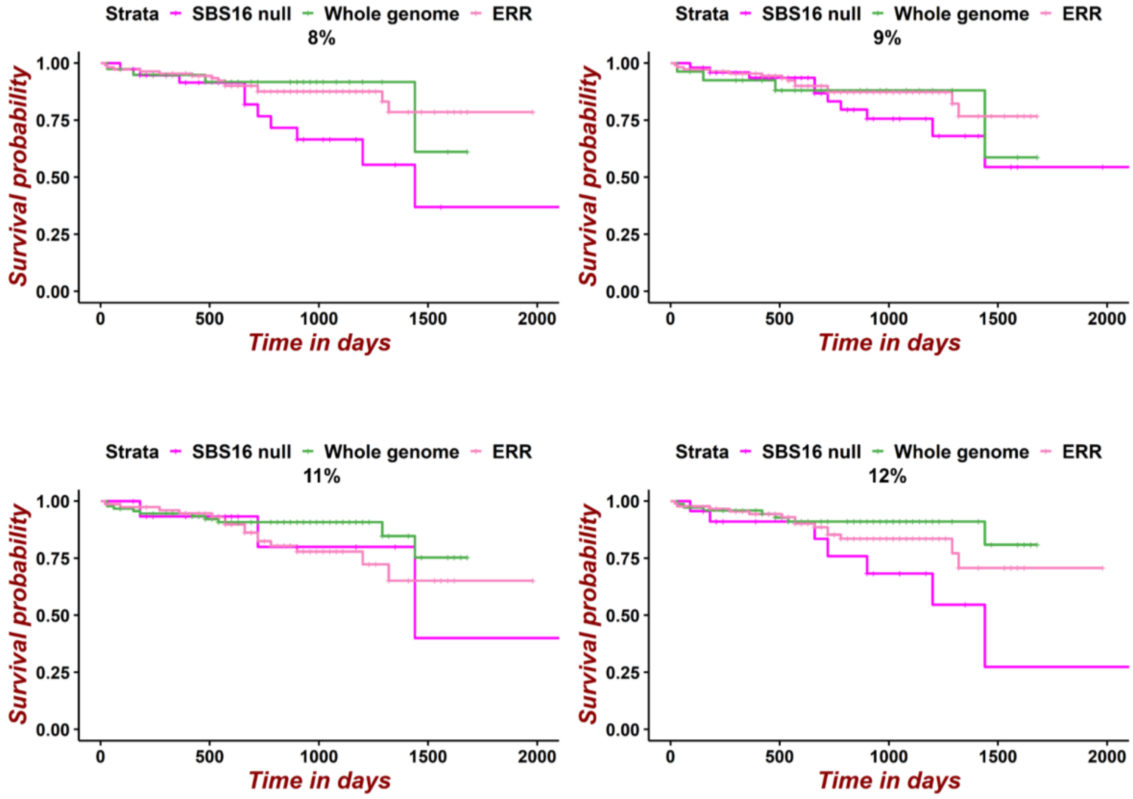


**Supplementary Figure 5 |** Survival analysis in LIRI-JP project, male patients, in different SBS16 thresholds from 8% to 12%. For each sample in each threshold, SBS16 was considered whole genome positive if found above the threshold in entire genome approach, ERR-only positive if found above the threshold in ERR-only approach, and negative otherwise.

### Supplementary Figure 6

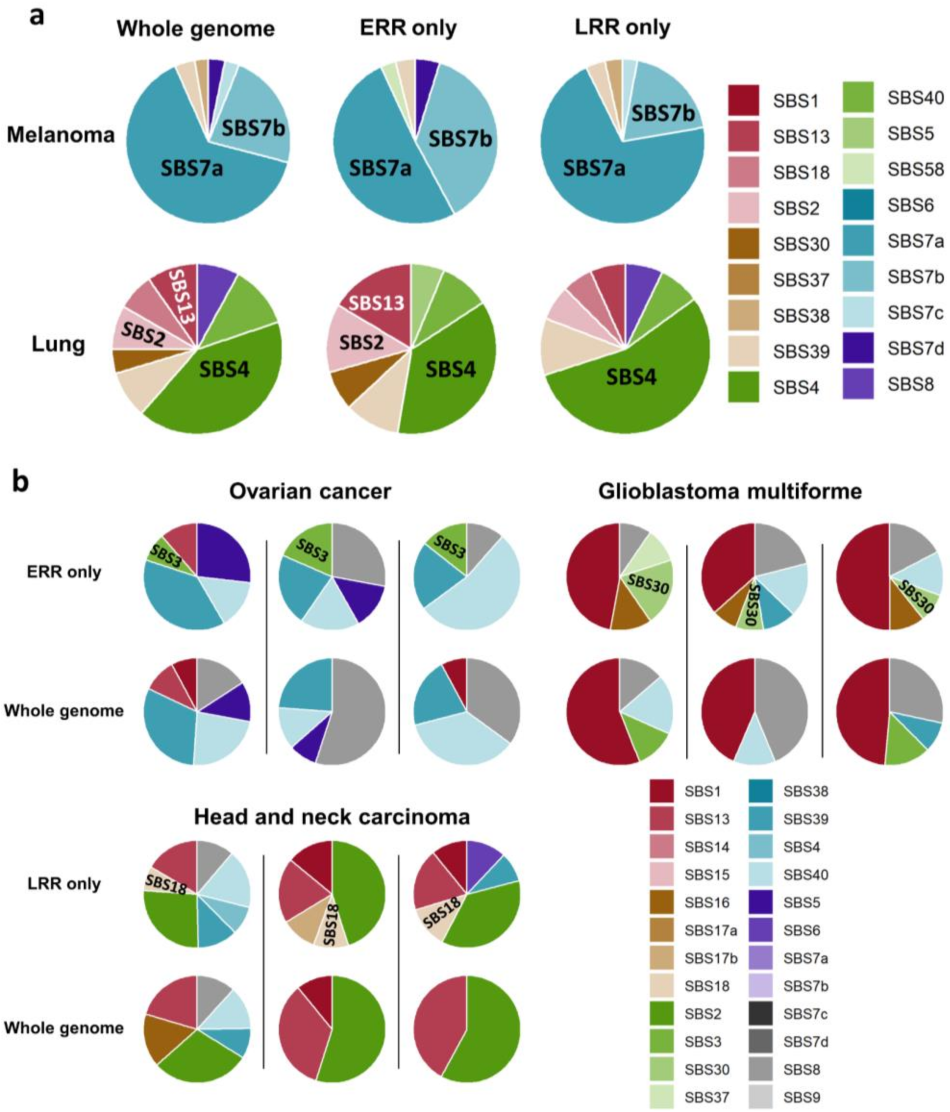


**Supplementary Figure 6 | a,** Pie charts illustrating the changes of SBS signatures' contribution in entire genome, ERR only, and LRR only, within specific projects. The average contribution of each signature in melanoma and in lung cancer samples is presented. **b,** Pie charts of individual samples' SBS signatures contribution for which specific signature is found only when using focused signature identification approach. Upper: SBS3 in three ovarian cancer samples (left) and SBS30 in three Glioblastoma multiforme samples (right), found only in ERR-focused approach. Lower: SBS18 in three head and neck carcinoma samples, found only in LRR-focused approach.

### Supplementary Figure 7

a

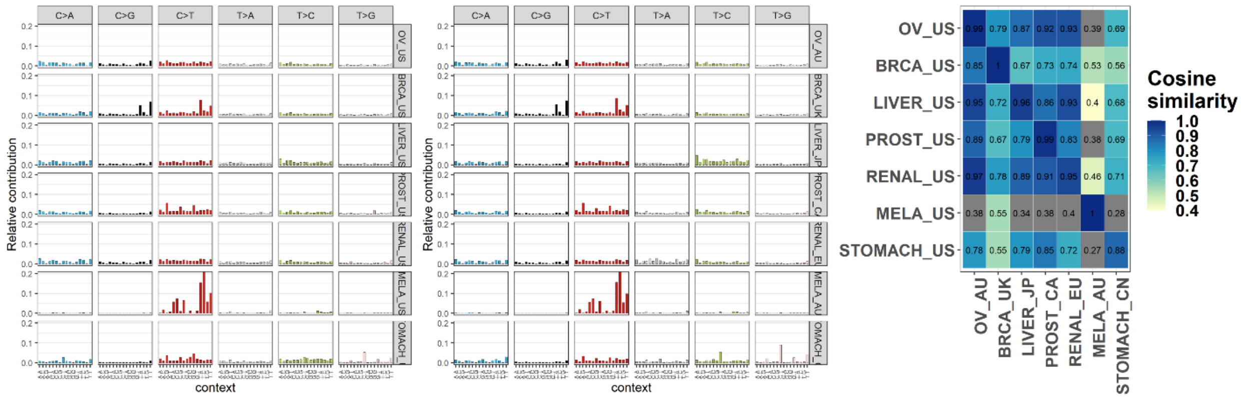


b

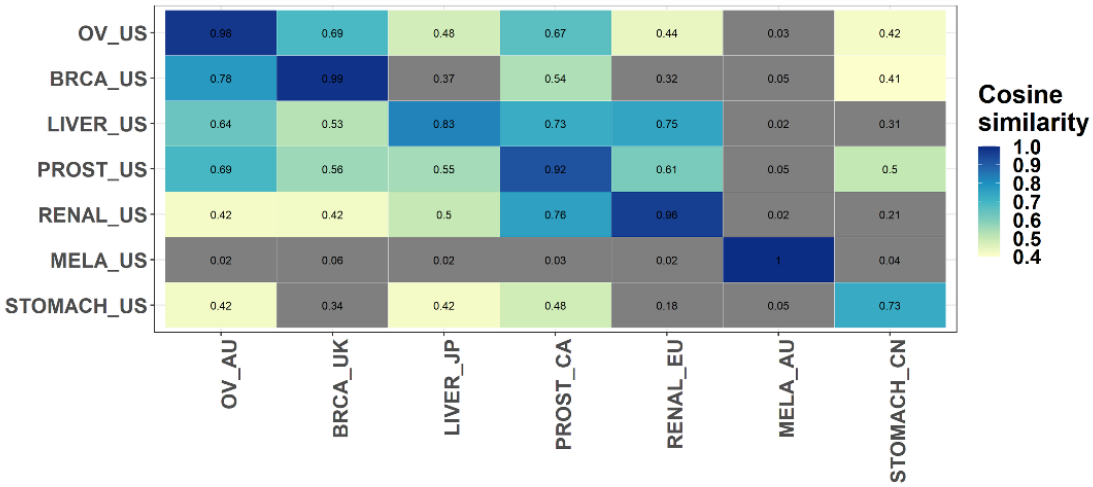


**Supplementary Figure 7 | a,** Mutational profiles similarity between TCGA and ICGC components of PCAWG. Seven cancer types published in both parts were inspected. Left: For each project from TCGA part, the mean mutation profile is presented; Middle: mutational profiles of the ICGC part projects; Right: correlation matrix of the mutational profiles. Note the high similarity along the diagonal line that represent the comparison of the same cancer type in the two projects. **b,** Similar analysis was performed in the signature level. Each project was represented by the relative contribution of each signature and the cosine similarities between the projects are shown.

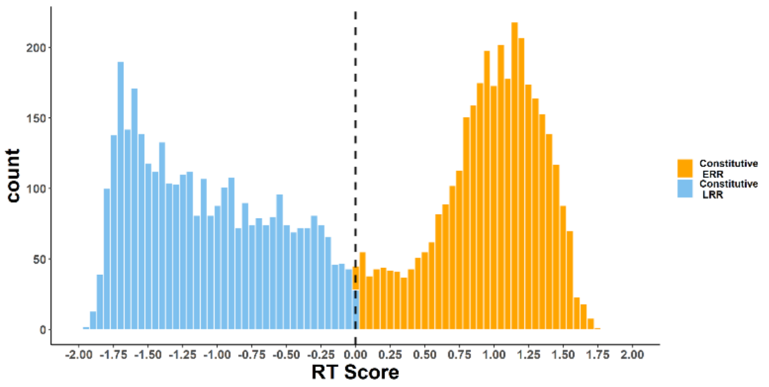


**Supplementary Figure 8 |** Histogram showing RT scores of constitutive RT regions. Values > 0 are considered as ERR (706 Mb), whereas values < 0 are considered as LRR (583 Mb).

### Supplementary Table 1

| Signature | Proposed etiology | Comment | RT Bias | Chromatin structure-independent RT bias | Cancer type (per COSMIC) |
|---|---|---|---|---|---|
| **SBS1** | Endogenous, clock-like, deamination of 5-methylcytosine to thymine | Found in most cancers and in normal cells, correlated with age | **late** | **late, only in active regions** | Pan-cancer |
| **SBS2** | AID/APOBEC cytidine deaminases | | **early** | **early, only in active regions** | Signature is pan-cancer, RT bias is cancer-type(s) dependant |
| **SBS3** | Defective HDR | | **early** | **early** | cancer type(s) dependant |
| **SBS4** | Direct DNA damage by tobacco smoke mutagens | | **late** | **late, only in inactive regions** | cancer type(s) dependant |
| **SBS5** | SBS5 is clock-like in that the number of mutations in most cancers and normal cells correlates with the age of the individual. | | **early** | **early** | Pan-cancer |
| **SBS7a** | Exposure to ultraviolet light | Possibly be the consequence of just one of the two major known UV photoproducts, cyclobutane pyrimidine dimers or 6-4 photoproducts | **late** | **late** | cancer type(s) dependant |
| **SBS7b** | Exposure to ultraviolet light | See SBS7a | **early** | **early** | cancer type(s) dependant |
| **SBS8** | Unknown | Recently shown to be due to late replications error in cancer | **late** | **late, only in inactive regions** | Pan-cancer |
| **SBS9** | May be due in part to mutations induced during replication by polymerase eta as part of somatic hypermutation in lymphoid cells. | Have elevated numbers of mutations attributed to SBS9 in IGHC-mutated CLL cells | **late** | **Not established** | cancer type(s) dependant |

| | | | | | |
|---|---|---|---|---|---|
| **SBS12** | Unknown | | **neutral** | **late, only in inactive regions** | cancer type(s) dependant |
| **SBS13** | AID/APOBEC cytidine deaminases | | **early** | **early, only in active regions** | Signature is pan-cancer, RT bias is cancer-type(s) dependant |
| **SBS16** | Unknown | Male-gender, alcohol and tobacco consumption associated. Contaminated by SBS5 and thus should handle carefully | **early** | **early** | cancer type(s) dependant |
| **SBS17a+b** | Unknown | | **late** | **Not established** | cancer type(s) dependant |
| **SBS18** | Possibly damage by reactive oxygen species. | | **late** | **Not established** | cancer type(s) dependant |
| **SBS39** | Unknown | | **early** | **neutral** | cancer type(s) dependant |
| **SBS40** | Unknown | Pan-cancer, highly common signature | **early** | **early** | Pan-cancer |

**Supplementary Table 1 |** SBS signatures with a clear RT bias (|delta|>0.05 in at least one project, and the relation between the RT bias and chromatin structure.

### Supplementary Table 2

| Cell line | Category | Base counts - Early | Base counts - Late |
|---|---|---|---|
| FM-55-P | Active | 877203579 | 113085620 |
| | Inactive | 340617436 | 981654018 |
| | Repressed | 240413947 | 148213676 |
| | Heterochromatin | 23096987 | 68541934 |
| BT-549 | Active | 949800000 | 161600000 |
| | Inactive | 432800000 | 865300000 |
| | Repressed | 303200000 | 147000000 |
| | Heterochromatin | 36400000 | 59200000 |
| HepG2 | Active | 860467000 | 133168000 |
| | Inactive | 199824000 | 1023739000 |
| | Repressed | 236067000 | 154291000 |
| | Heterochromatin | 22451000 | 68658000 |
| PANC-1 | Active | 914900000 | 196500000 |
| | Inactive | 309500000 | 988600000 |
| | Repressed | 320200000 | 130000000 |
| | Heterochromatin | 48300000 | 47300000 |
| A549 | Active | 935700000 | 158400000 |
| | Inactive | 239600000 | 1004000000 |
| | Repressed | 307700000 | 131000000 |
| | Heterochromatin | 24900000 | 61000000 |

**Supplementary Table 2 | Replication timing and chromatin accessibility regions.** Four chromatin states (Active, Inactive, Repressed and Heterochromatin) intersected with replication timing regions (Early, Late) in 5 cell lines' replication timing regions.
